# Fibroblast growth factor-9 expression in airway epithelial cells amplifies the type I interferon response and alters influenza A virus pathogenesis

**DOI:** 10.1101/2021.12.22.473953

**Authors:** Bradley E. Hiller, Yongjun Yin, Yi-Chieh Perng, Ítalo de Araujo Castro, Lindsey E. Fox, Marissa C. Locke, Kristen J. Monte, Carolina B. López, David M. Ornitz, Deborah J. Lenschow

## Abstract

Influenza A virus (IAV) preferentially infects conducting airway and alveolar epithelial cells in the lung. The outcome of these infections is impacted by the host response, including the production of various cytokines, chemokines, and growth factors. Fibroblast growth factor-9 (FGF9) is required for lung development, can display antiviral activity *in vitro*, and is upregulated in asymptomatic patients during early IAV infection. We therefore hypothesized that FGF9 would protect the lungs from respiratory virus infection and evaluated IAV pathogenesis in mice that overexpress FGF9 in club cells in the conducting airway epithelium (FGF9-OE mice). However, we found that FGF9-OE mice were highly susceptible to IAV and Sendai virus infection compared to control mice. FGF9-OE mice displayed elevated and persistent viral loads, increased expression of cytokines and chemokines, and increased numbers of infiltrating immune cells as early as 1 day post-infection (dpi). Gene expression analysis showed an elevated type I interferon (IFN) signature in the conducting airway epithelium and analysis of IAV tropism uncovered a dramatic shift in infection from the conducting airway epithelium to the alveolar epithelium in FGF9-OE lungs. These results demonstrate that FGF9 signaling primes the conducting airway epithelium to rapidly induce a localized, protective IFN and proinflammatory cytokine response during viral infection. Although this response protects the airway epithelial cells from IAV infection, it allows for early and enhanced infection of the alveolar epithelium, ultimately leading to increased morbidity and mortality. Our study illuminates a novel role for FGF9 in regulating respiratory virus infection and pathogenesis.

**Author Summary:** Influenza viruses are respiratory viruses that cause significant morbidity and mortality worldwide. In the lungs, influenza A virus primarily infects epithelial cells that line the conducting airways and alveoli. Fibroblast growth factor-9 (FGF9) is a growth factor that has been shown to have antiviral activity and is upregulated during early IAV infection in asymptomatic patients, leading us to hypothesize that FGF9 would protect the lung epithelium from IAV infection. However, mice that express and secrete FGF9 from club cells in the conducting airway had more severe respiratory virus infection and a hyperactive inflammatory immune response as early as 1 day post-infection. Analysis of the FGF9-expressing airway epithelial cells found an elevated antiviral and inflammatory interferon signature, which protected these cells from severe IAV infection. However, heightened infection of alveolar cells resulted in excessive inflammation in the alveoli, resulting in more severe disease and death. Our study identifies a novel antiviral and inflammatory role for FGFs in the lung airway epithelium and confirms that early and robust IAV infection of alveolar cells results in more severe disease.

## Introduction

Influenza A virus (IAV) is a respiratory pathogen in the *Orthomyxoviridae* family that causes both devastating pandemics and seasonal epidemics, resulting in significant morbidity and mortality worldwide. Once inhaled, IAV primarily targets differentiated epithelial cells of the upper and lower respiratory tract, including ciliated cells, club cells, and type I and type II alveolar epithelial cells (AT1 and AT2 cells, respectively)^1^. Infection of these cells, in addition to abortive infection of immune cells, leads to virus recognition by multiple pattern recognition receptors (PRRs), resulting in the production of proinflammatory cytokines and type I interferons (IFNs) which activate hundreds of IFN-stimulated genes (ISGs)^2–4^. While antiviral ISGs reduce viral burden and protect neighboring cells from IAV infection, excessive inflammation from type I IFN-induced cytokines and chemokines can exacerbate lung disease. Therefore, proper resolution of IAV infection requires a coordinated respiratory epithelial and immune cell response to limit viral replication, contain inflammation, and promote repair of the lung epithelium.

Multiple secreted proteins in the lung, including cytokines and growth factors, can regulate virus infection, the inflammatory response, and post-infection repair to combat severe IAV disease. Fibroblast growth factors (FGFs) are a family of secreted proteins that regulate tissue maintenance, metabolism, regeneration, and repair of adult tissues by binding to the extracellular domain of four transmembrane tyrosine kinase FGF receptors (FGFRs)^5, 6^. In the developing lung, FGF1, FGF2, FGF3, FGF7, FGF9, FGF10, and FGF18 all perform essential regulatory functions mediating organogenesis^7–12^. FGF signaling is required for epithelial and mesenchymal maintenance, and elevated FGF signaling or FGF expression can have beneficial or detrimental impacts upon lung biology. In the adult lung, increased expression of FGF2, FGF7, and FGF10 promotes repair after lung injury, and FGF10 also promotes resolution of acute lung injury and acute respiratory distress syndrome^13–15^. In contrast, FGF9 is elevated in tissue samples from patients with mild to severe idiopathic pulmonary fibrosis (IPF), and FGF9 and FGF18 both contribute to the development of IPF in tissue culture^16, 17^.

FGFs in the lung have diverse roles during respiratory virus infection. During IAV infection in mice, FGF7 increases the severity of IAV infection by accelerating AT2 cell infection and proliferation^18^. During IAV infection of both humans and mice, FGF2 expression is elevated in the lungs^19^. FGF2 administration in mice reduced the severity of IAV disease while knockout of *Fgf2* increased IAV lung injury by impairing neutrophil recruitment and activation^19^. FGF10, however, protects the lung from severe IAV infection by driving epithelial repair^20^. During IAV infection of humans, increased FGF9 transcripts specifically in the serum of asymptomatic patients but not symptomatic patients has been observed at 1 dpi, suggesting that FGF9 may function in the early response to IAV infection^21^. In addition, a screen of 756 human secreted proteins identified members of the FGF9 subfamily (FGFs 9, 16, and 20) as inhibitors of vesicular stomatitis virus (VSV) replication^22^. Given these findings, we hypothesized that FGF9 may provide protective or antiviral functions during IAV infection in the adult lung.

To test our hypothesis, *Fgf9* was placed under doxycycline-inducible control directed by the *Scgb1a1* (*Cc10*, *Ccsp*) promoter, allowing us to temporally regulate FGF9 expression and secretion from club cells^23, 24^. Since club cells comprise the majority of the conducting airway epithelium in the mouse lung, we expected that expressing FGF9 in club cells would protect the entire conducting airway epithelium from IAV infection. However, FGF9-overexpressing mice displayed excessive lung inflammation and alveolar edema, increased cytokine and chemokine expression, accelerated infection in the alveolar space, and displayed increased mortality. Characterization of the conducting airway epithelium revealed that FGF9 overexpression primed the epithelium for an amplified type I IFN response that protected it from IAV infection. These findings provide important insights into how FGF signaling in the lung impacts the pathogenesis of respiratory virus disease.

## Results

### Generation and characterization of FGF9-overexpressing transgenic mice

To express *Fgf9* from club cells in the conducting airway epithelium with temporal specificity, TRE-Fgf9-IRES-eGfp transgenic mice^10^ were mated to transgenic mice with a reverse tetracycline transactivator (rtTA) driven under the *Scgb1a1* promoter (Scgb1a1-rtTA). This cross, hereafter referred to as FGF9-overexpressing (FGF9-OE) mice, results in the induction of the rtTA element primarily in club cells after replacing the standard mouse diet with doxycycline-containing chow (DOX) (**Fig. 1A**). Elevated levels of *Fgf9* RNA were detected by real-time quantitative PCR (RT-qPCR) from whole lungs of FGF9-OE mice upon DOX administration for 24, 48, and 72 hours compared to uninduced FGF9-OE mice (**Fig. 1B**). To further evaluate the induction of the TRE-Fgf9-IRES-eGfp cassette, lung sections isolated from FGF9-OE mice and mice containing only the TRE-Fgf9-IRES-eGfp or Scgb1a1-rtTA transgene (hereafter referred to as littermate “control” mice) were analyzed after 3 days of DOX administration. Lung sections from DOX-treated control mice or from uninduced FGF9-OE mice displayed no detectable eGFP activity in the airway epithelium (**Fig. 1C**, **Supp. Fig. 2**). In contrast, DOX-treated FGF9-OE mice showed eGFP fluorescence co-localized within the club cells as denoted by SCGB1A1 staining. Furthermore, histological analysis of the lung sections revealed no gross differences in the conducting airway or alveolar epithelium or evidence of cellular infiltrates in the lungs between DOX-treated FGF9-OE mice, DOX-treated control mice, or FGF9-OE mice without DOX (**Fig. 1D**). To confirm whether 3 days of FGF9 overexpression altered epithelial cell number in the lungs, FGF9-OE and control mice were administered DOX for 3 days, and whole, right lung lobes were harvested and analyzed by flow cytometry (**Supp. Fig. 1A**). Consistent with our histological analysis of lung sections, we detected no differences in the number of total epithelial cells (EpCAM^+^), conducting airway epithelial cells (EpCAM^+^CD24^+^) or alveolar epithelial cells (EpCAM^+^CD24^−^) between control and FGF9-OE mice (**Fig. 1E**).

**Figure 1.**
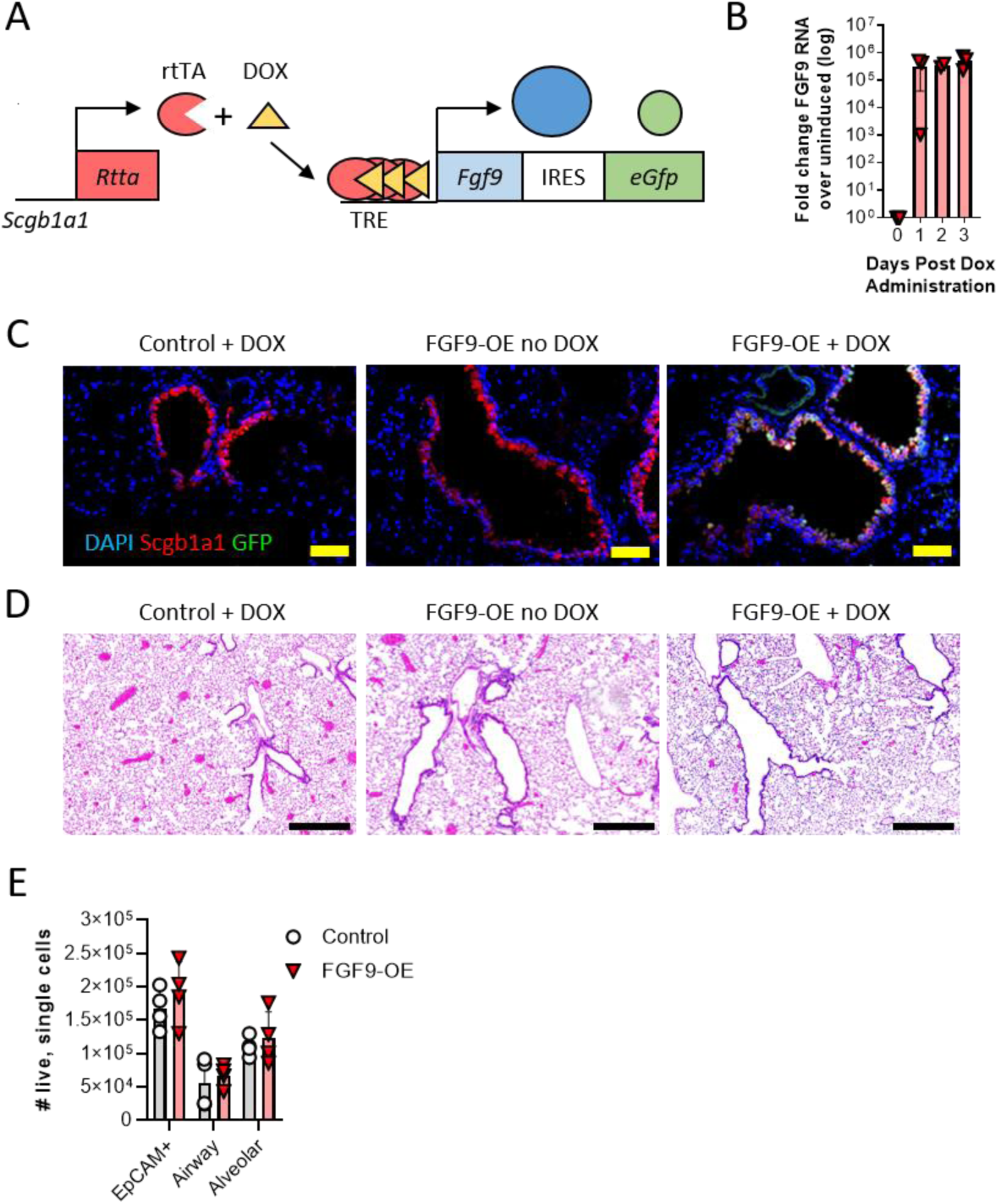
FGF9 overexpression from club cells after 3 days of DOX administration does not alter lung histology. (A) Schematic of the doxycycline chow (DOX)-inducible double-transgenic mouse breeding resulting in single-transgenic control mice (Scgb1a1-rtTA or TRE-Fgf9-IRES-eGfp) or double-transgenic FGF9-OE mice (Scgb1a1-rtTA and TRE-Fgf9-IRES-eGfp). (B) DOX was administered to FGF9-OE mice for 1-3 days and *Fgf9* was measured in whole lung RNA by RT-qPCR. Data are represented as fold change of *Fgf9* compared to the uninduced lung (0 days post DOX administration). (C-D) Representative images of control and FGF9-OE mouse lung sections after 3 days of DOX administration; (C) stained with DAPI (blue), anti-SCGB1A1 (red), and eGFP (green), scale bars = 50 µm, (D) stained with H&E, scale bars = 500 µm. (E) FGF9-OE and control mice were given DOX for 3 days, lungs were harvested, and single cell suspensions were analyzed for epithelial cells by flow cytometry as described in materials and methods. Data are represented as mean ± SEM and analyzed by unpaired student’s *t* test.

### FGF9-OE mice are more susceptible to respiratory virus infection than control mice

Given the ability of FGF9 to inhibit recombinant VSVs^22^, we hypothesized that FGF9 may inhibit IAV infection and thereby protect the lungs from severe disease. To test this hypothesis, FGF9-OE and littermate control mice were continuously administered DOX beginning 3 days prior to infection (d-3), infected intranasally (i.n.) (d0) with IAV A/WSN/33 (WSN), and then monitored for weight loss and survival. Although we expected the FGF9-OE mice to be protected from IAV infection, all FGF9-OE mice lost weight and succumbed to infection between 5-9 dpi (**Fig. 2A-B**). In contrast, while the control mice initially lost weight at the same rate as the FGF9-OE mice, 100% of the control mice recovered and survived WSN infection.

**Figure 2.**
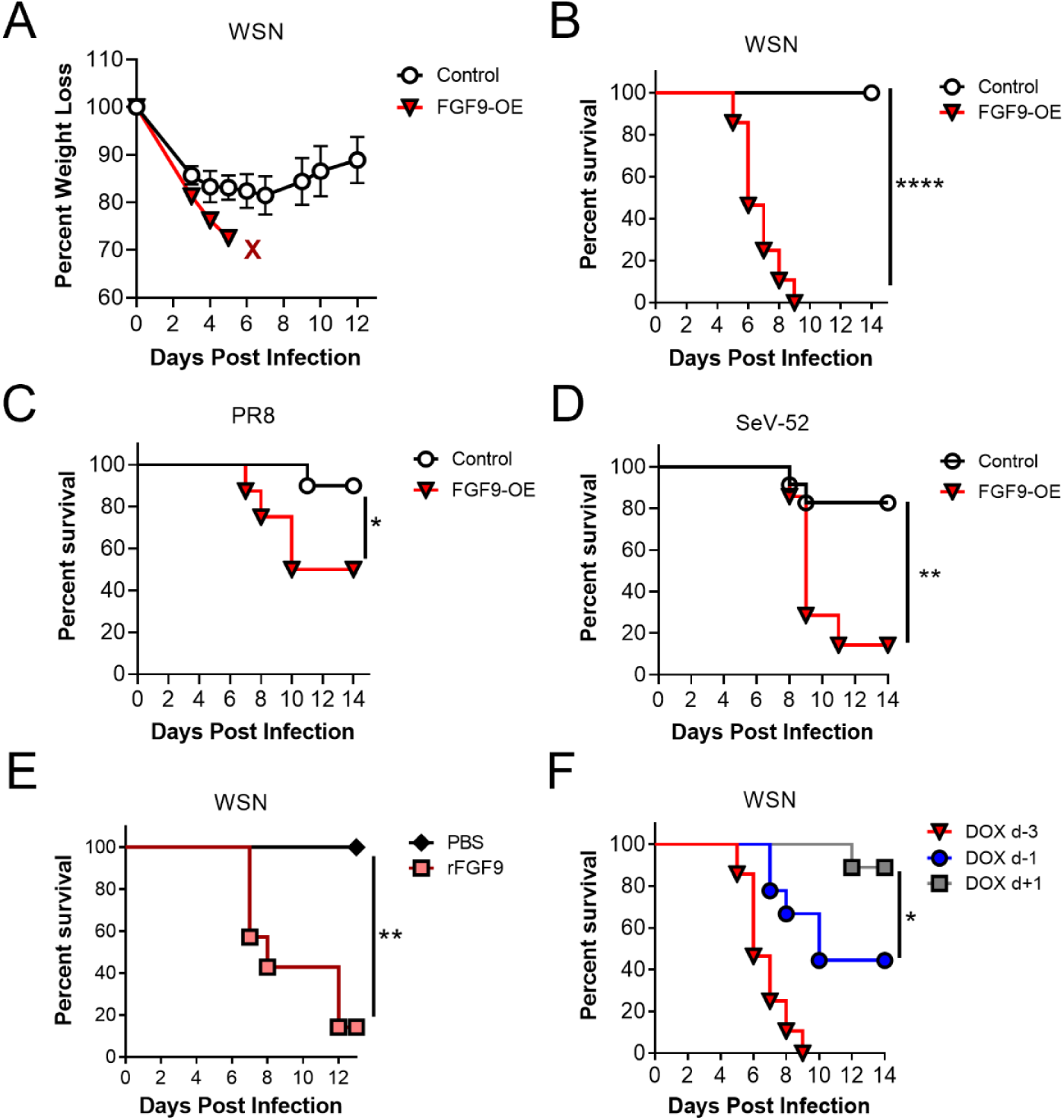
FGF9-OE mice are more susceptible to respiratory virus infection than control mice. (A-B) FGF9-OE and control mice were administered DOX beginning 3 days prior to infection (d-3), inoculated intranasally (i.n.) on d0 with 6×10^4^ PFU WSN, and monitored for (A) weight loss (control n=12, FGF9-OE n=8) and (B) survival (control n=31, FGF9-OE n=28). (C-D) DOX- induced FGF9-OE and control mice were inoculated i.n. on d0 with 5 PFU PR8 (control n=10, FGF9-OE n=8) (C) or 1×10^4^ PFU SeV-52 (control n=23, FGF9-OE n=7) (D) and were monitored for survival. (E) WT FVB/NJ mice were treated i.n. with PBS (n=7) or 5 µg rFGF9 (n=7) on d-3 and again concurrently with WSN infection on d0 and were monitored for survival. (F) Survival curve of WSN-infected FGF9-OE mice treated with DOX 1 day pre-infection (d-1, n=9) or 1 dpi (d+1, n=9) compared to historic d-3 DOX administration from panel B. For all experiments, weight loss and survival were monitored until 14 dpi. Data were pooled from 2 or more separate experiments. Data are represented as mean ± SEM and were analyzed by Mantel-Cox test (*, p < 0.05; **, p < 0.01; ****, p < 0.0001).

To determine whether FGF9-OE mice were similarly more susceptible to other respiratory viruses, we evaluated survival of DOX-treated mice infected with IAV A/PR/8/34 (PR8) or Sendai virus (SeV-52). While the majority of control mice recovered from PR8 infection (**Fig. 2C**) and SeV-52 infection (**Fig. 2D**), the FGF9-OE mice were more susceptible to both respiratory viruses compared to control mice. Together, these results demonstrate that overexpression of FGF9 from club cells leads to increased susceptibility to respiratory virus infection. To characterize this FGF9-driven increase in disease, we focused on elucidating the mechanisms of heightened WSN pathogenesis in FGF9-OE mice since the difference in survival between FGF9-OE and control mice was most striking during WSN infection.

Because club cells in the wild-type adult mouse lung do not specifically express high levels of FGF9, we wanted to dispel the possibility that the FGF9-OE mice are more susceptible to WSN infection due to unintended effects of the double-transgenic mouse model unrelated to FGF9 expression. Therefore, we next determined whether recombinant FGF9 (rFGF9) similarly promoted susceptibility to WSN infection. WT FVB/NJ mice were treated with 5 µg rFGF9 or PBS i.n. on d-3 and again concurrently with WSN infection on d0, and survival was monitored. While all PBS-treated mice recovered from WSN infection, nearly all rFGF9-treated mice succumbed to the infection (**Fig. 2E**), confirming that increased FGF9 in the respiratory tract enhanced influenza severity. Next, to determine when FGF9 overexpression was required for increased susceptibility to IAV infection, we induced FGF9 overexpression at varying time points. FGF9-OE mice were given DOX 1 day prior to infection (d-1) or 1 day post infection (d+1), were infected with WSN at d0, and were monitored for survival. FGF9-OE mice given DOX on d-1 displayed intermediate lethality compared to what we observed when treatment was initiated on day -3, with approximately 45% of the mice recovering from infection (**Fig. 2F**). However, if DOX administration was started on d+1, then nearly all the mice survived WSN infection. Taken altogether, these results demonstrate that forced overexpression of FGF9 in club cells increases susceptibility to different respiratory viruses, but only if overexpression is initiated prior to infection.

### FGF9-OE mice have delayed viral clearance and exacerbated inflammation during IAV infection

Increased susceptibility to WSN required FGF9 overexpression prior to infection. However, lethality in FGF9-OE mice did not occur until 5-6 dpi, likely due to increased viral burden, altered viral clearance, or an excessive inflammatory response. First, to evaluate the impact of FGF9 overexpression on viral burden, the lungs of DOX-treated, WSN-infected FGF9-OE and control mice were harvested at 1, 3, and 6 dpi to measure infectious virus by plaque assay. At 1 dpi, the levels of replicating virus did not differ between the control and FGF9-OE mice (**Fig. 3A**). By 3 dpi, the FGF9-OE mice sustained viral titers of 10^5^ PFU/ml while the viral titers in control mice were beginning to decline, with an approximate 1-log reduction in viral burden. At 6 dpi, nearly all the control mice had cleared replicating virus from their lungs, while the FGF9-OE mice failed to clear infectious virus and still displayed titers as high as 10^4^ PFU/ml.

**Figure 3.**
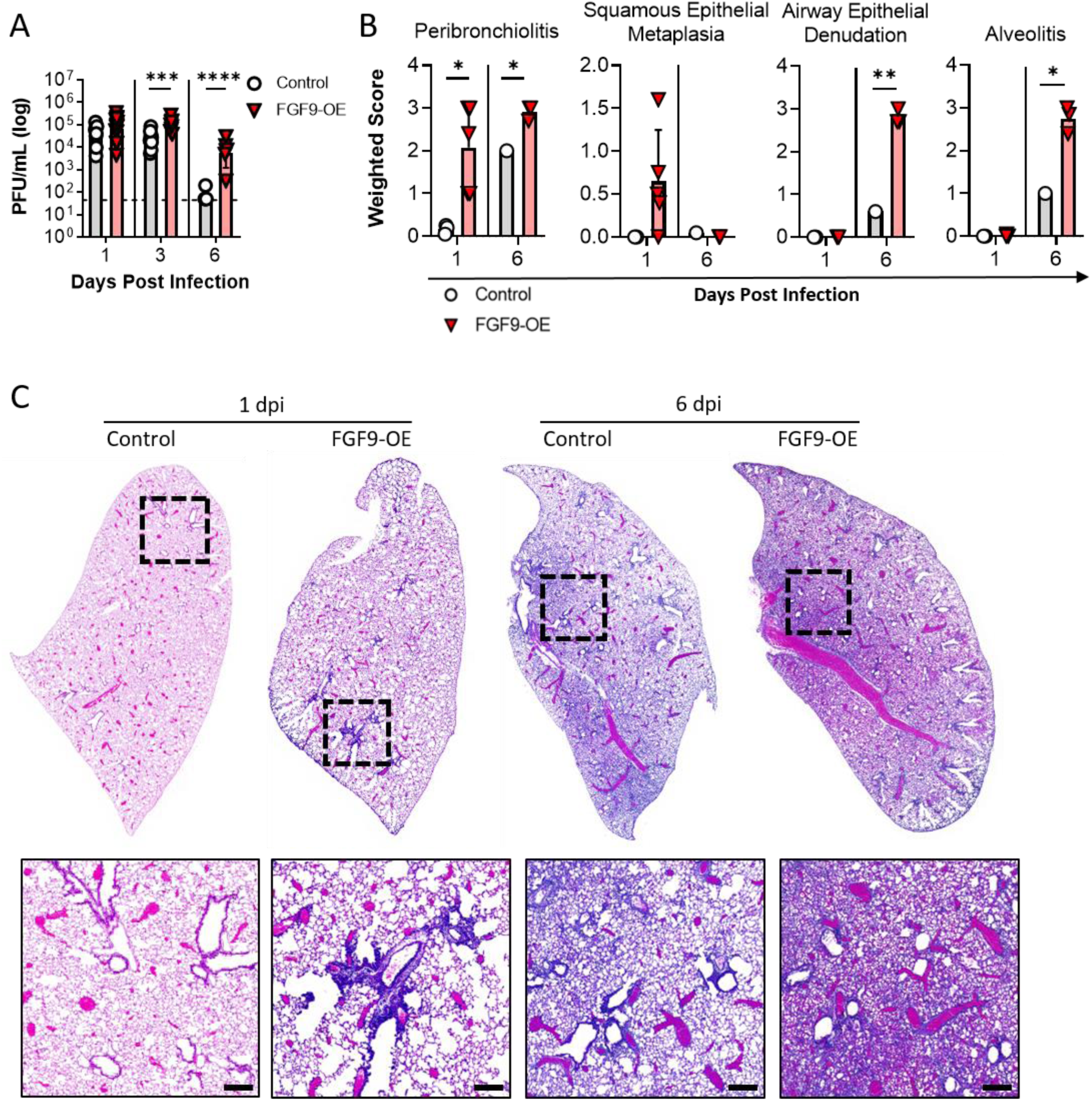
FGF9-OE mice sustain higher viral titers and have increased lung inflammation. (A-C) FGF9-OE and control mice were administered DOX beginning on d-3 and infected on d0 with 6×10^4^ PFU WSN i.n. Lungs were harvested at 1, 3, and 6 dpi. At each time point, n=6-13 control or FGF9-OE mice were analyzed. (A) Infectious virus was quantified by plaque assay; dotted line represents 50 PFU/ml (limit of detection). (B) H&E-stained slides at 1 dpi (control n=3, FGF9-OE n=5) and 6 dpi (control n=1, FGF9-OE n=3) were scored for peribronchiolitis, squamous epithelial metaplasia, airway epithelial denudation, and alveolitis as described in the Materials and Methods. (C) Representative images of H&E-stained control and FGF9-OE whole lung sections at 1 and 6 dpi; (bottom) corresponding magnified insets (scale bars = 200 µm). Data are represented as mean ± SEM and were analyzed within each time point by unpaired student’s *t* test (*, p < 0.05; **, p < 0.01; ***, p < 0.001; ****, p < 0.0001).

We next scored histopathology in tissue sections collected from DOX-treated FGF9-OE and control mice at early (1 dpi) and late (6 dpi) infection time points. At 1 dpi, we observed thickening of the FGF9-OE airways driven by significantly increased peribronchiolitis and a higher prevalence of squamous epithelial cell metaplasia (**Fig. 3B-C**), compared to relatively low levels of peribronchiolitis and a lack of squamous epithelial cell metaplasia in control lung sections. By 6 dpi, control mouse lung sections displayed increased peribronchiolitis and alveolitis characterized by a moderate number of infiltrating leukocytes in the alveolar space. In contrast, we observed significantly more severe alveolitis in FGF9-OE lung sections at 6 dpi, characterized by a high number of infiltrating leukocytes, alveolar edema, and alveolar consolidation. We additionally observed significant denuding of the FGF9-OE airway epithelium at 6 dpi with severe peribronchiolitis around the remaining conducting airways (**Fig. 3B-C**).

Altogether, these results demonstrate that club cell-driven FGF9 overexpression led to elevated and sustained viral titers in the lung tissue. We also observed pervasive inflammation of the airway epithelium and, most significantly, the lung parenchyma during IAV infection, correlating to when the FGF9-OE mice began to succumb to infection.

### FGF9-OE mice display an accelerated and elevated inflammatory response to IAV

Respiratory epithelial cells are the primary target of IAV replication and are important for the initial amplification of cytokine and chemokine expression^25^. Multiple experimental studies have shown that excessive cytokine and chemokine expression during IAV infection correlates with increased tissue damage and severity of disease^26–28^. Given the increased alveolar inflammation we observed in FGF9-OE mouse lungs (**Fig. 3B-C**), we next evaluated if these mice displayed elevated cytokine and chemokine expression throughout infection. FGF9-OE and control mice were administered DOX beginning at d-3, infected with WSN at day 0, and the right lung lobes were taken from mice after 3 days of DOX but without infection (0 dpi) or at 1, 3, or 5 dpi (**Fig. 4A**). Cytokine and chemokine protein levels were then quantified in lung homogenates by multiplex array analysis. In mice given DOX only (0 dpi), several analytes were statistically increased in FGF9-OE lungs compared to control lungs, including IL-7, IL-9, CCL2, CCL4, CXCL9, and CXCL10, although none of these analytes were increased above 2-fold (**Fig. 4B**). Strikingly, 16 cytokines and chemokines were significantly elevated in lungs isolated from the FGF9-OE mice at 1 dpi compared to control mice, including G-CSF, TNFα, IFNγ, IL-6, CCL2, CCL4, and CXCL9 (**Fig. 4B-C**). Multiple cytokines and chemokines remained highly expressed at 3 and 5 dpi in the FGF9-OE lysates, including G-CSF, TNFα, IL-6, and CCL2 (**Fig. 4B**).

**Figure 4.**
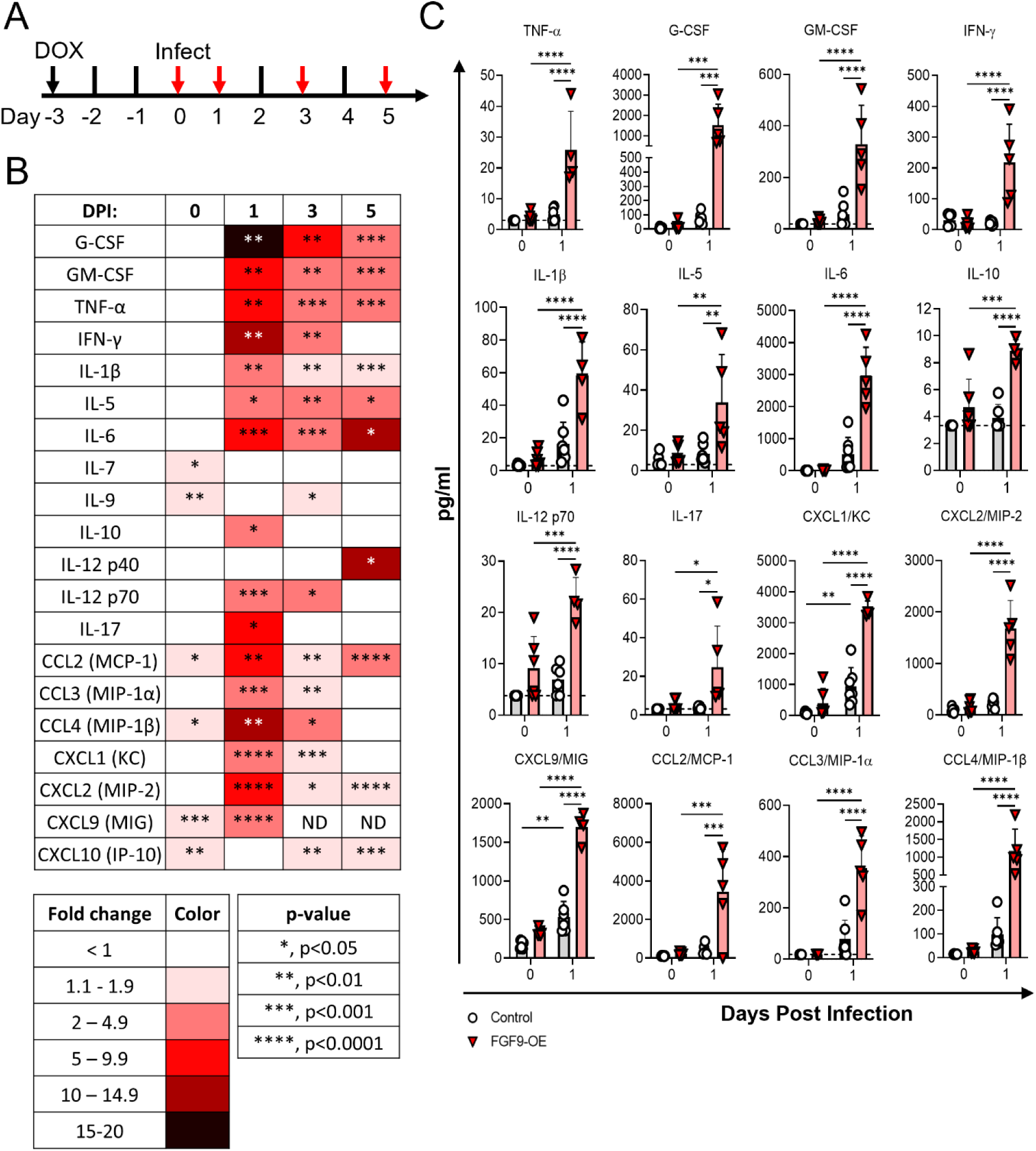
Cytokine and chemokine expression is elevated in FGF9-OE lungs during IAV infection. (A-C) FGF9-OE and control mice were administered DOX beginning on d-3 and infected on d0 with 6×10^4^ PFU WSN i.n. (A) Lungs were harvested prior to infection on d0 (0 dpi) and at 1, 3, and 5 dpi (red arrows) and whole lung homogenates were analyzed by multiplex cytokine and chemokine analysis. (B) Data represented in a table as mean fold change of FGF9-OE samples over control samples for each analyte at 0, 1, 3, and 5 dpi. Data statistics were analyzed within each time point using unpaired student’s *t* test (*, p < 0.05; **, p < 0.01; ***, p < 0.001; ****, p < 0.0001). No symbol indicates no significant difference in analyte expression between FGF9-OE and control samples; ND, not determined. (C) Fold change of 16 selected analytes significantly increased in FGF9-OE samples at 1 dpi compared to control samples, represented as mean ± SEM at 0 and 1 dpi. Data were analyzed across time points by two-way ANOVA (*, p < 0.05; **, p < 0.01; ***, p < 0.001; ****, p < 0.0001). Dotted lines represent limit of detection. At each time point, n=5-7 control or FGF9-OE mice were analyzed.

The initial surfeit of cytokines and chemokines we observed in the FGF9-OE lung lysates at 1 dpi suggests that FGF9 had driven an increased inflammatory response to infection considering that there were no significant differences in viral titers at 1 dpi between FGF9-OE and control mice (**Fig. 3A**). Since we detected minor increases in several cytokines and chemokines with just 3 days of FGF9 overexpression in the absence of infection (**Fig. 4B-C**), we wanted to further characterize the immune environment of the FGF9-OE and control lungs both without WSN infection and at 1 dpi by analyzing conventional immune cell populations. Single-cell suspensions from whole, right lungs of DOX-induced FGF9-OE and control mice were generated after 3 days of DOX but without infection (0 dpi) and at 1 dpi, and the total number of innate immune cells were quantified by flow cytometry (**Supp. Fig. 1B**). While there were no significant differences in the total number of CD45^+^ cells or any immune cell subtypes measured at 0 dpi, FGF9-OE lungs displayed a 2-fold increase in the total number of CD45^+^ cells at 1 dpi (**Fig. 5A**). FGF9-OE lungs had slightly elevated (< 2-fold) numbers of alveolar macrophages (**Fig. 5B**) and eosinophils (**Fig. 5C**) at 1 dpi compared to control lungs, as well as a 7-fold increase in neutrophils (**Fig. 5D**), a 2-fold increase in the number of CD103^+^ DCs (**Fig. 5E**) and CD11b^+^ DCs (**Fig. 5F**), a 3-fold increase in the number of Ly6C^+^ monocytes (**Fig. 5G**) and CD11b^+^ macrophages (**Fig. 5H**), and a trend toward elevated moDC numbers (**Fig. 5I**). Taken together, these data indicate that, while 3 days of FGF9 overexpression results in minor increases in select cytokines and chemokines, the additional stimulus of infection is required to elicit hyperactive expression of cytokines and chemokines and increased immune cells as early as 1 dpi in FGF9-OE lungs compared to control lungs.

**Figure 5.**
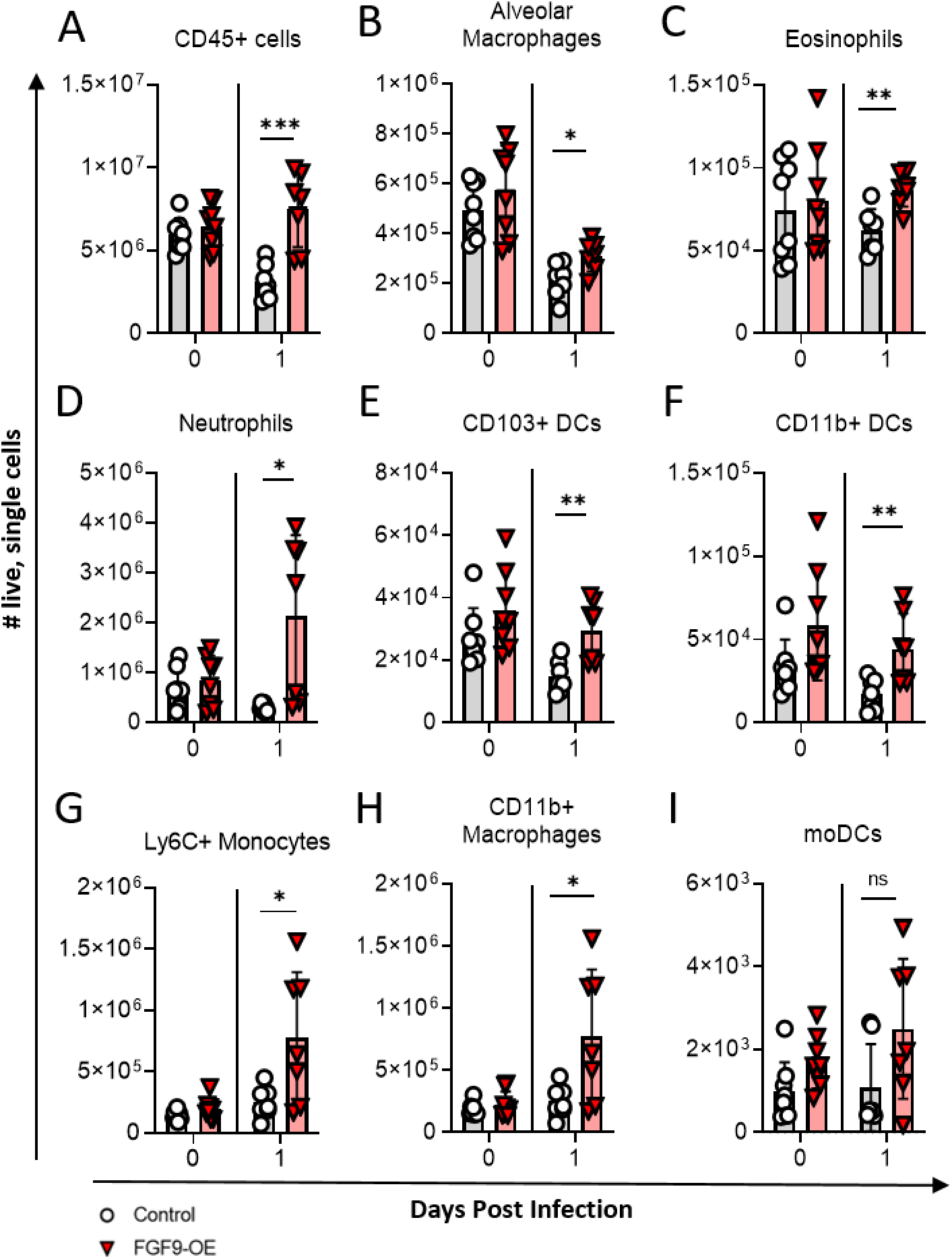
Increased numbers of innate immune cells infiltrate into FGF9-OE lungs at 1 dpi. (A-I) FGF9-OE and control mice were administered DOX beginning on d-3 and infected on d0 with 6×10^4^ PFU WSN i.n. Lung single cell suspensions were generated prior to infection (0 dpi) and at 1 dpi, and the total number of live cells was determined for the indicated immune cell subpopulation as described in materials and methods. Data are represented as mean ± SEM and analyzed within each time point by unpaired student’s *t* test (*, p < 0.05; **, p < 0.01; ***, p < 0.001). At each time point, n=7-8 control or FGF9-OE mice were analyzed.

### FGF9-OE airway epithelial cells have a strong IFN signature at 1 dpi

Secreted FGFs tightly associate with heparan sulfate proteoglycans on the surface of cells, which limits their diffusion and allows for robust autocrine and paracrine signaling on nearby neighboring cells^29–32^. In our FGF9-OE mouse model, FGF9 is secreted from club cells, which comprise the majority of the conducting airway epithelium in the mouse lung^8^. Therefore, we reason that the club cells, ciliated cells, and other epithelial cell types of the conducting airway are the cells that receive the most FGF9 stimulus and becoming transcriptionally altered.

To investigate this further, we sorted airway epithelial cells (CD45^−^CD326^+^CD24^+^) (**Supp. Fig. 1A**) from 1 dpi lung digests of DOX-induced FGF9-OE and control mice and performed RNA-seq on extracted RNA. Gene set enrichment analysis (GSEA) of the Gene Ontology Biological Processes database revealed that the top 10 most significant positively-enriched pathways in the FGF9-OE airway epithelial cells compared to control airway epithelial cells at 1 dpi were all related to the innate immune response and IFN signaling, including “Cellular response to IFNβ,” “Defense response to virus,” and “Inflammatory response” (**Fig. 6A**, **Supp. Fig. 3**, **Supp. Table 1**). *Ifnb1*, multiple *Ifna* subtypes (e.g. *Ifna11* and *Ifna14*), and *Ifnl2* were all upregulated by 1-3 log fold change in the FGF9-OE airway epithelial cells at 1 dpi, as well as multiple cytokines and chemokines (e.g. *Il6*, *Il33*, *Ccl9*, and *Cxcl11*), and known antiviral ISGs (e.g. *Isg15*, *Oas3*, and *Rsad2*) (**Fig. 6B**). Importantly, FGF9 was highly upregulated in the FGF9-OE airway epithelial cells, but was lowly-expressed in the control airway epithelial cells. RT-qPCR analysis of RNA extracted from sorted airway epithelial cells at 1 dpi validated the upregulation of several of these genes, including *Ifnb1*, *Isg15*, and *Rsad2* (viperin) in FGF9-OE airway epithelial cells as compared to cells isolated from control mice (**Fig. 6C**). We also assessed ISG15 protein expression by immunofluorescence microscopy. At 1 dpi we saw a modest upregulation of ISG15 protein expression in the control airway epithelium, but in the FGF9-OE lungs, ISG15 expression was dramatically upregulated in the airway epithelial cells (**Fig. 6D**). Finally, to determine if FGF9 signaling alone resulted in this increased IFN signature in the conducting airway epithelium, we analyzed expression of several genes in FGF9-OE mice that were treated with DOX for 3 days but were not infected (0 dpi). RT-qPCR analysis of sorted airway epithelial cells from FGF9-OE and control mice at 0 dpi revealed no significant differences in *Ifnb1*, *Isg15*, and *Rsad2* expression (**Fig. 6C**). We also observed no ISG15 protein expression by immunofluorescence microscopy in the lungs of FGF9-OE mice following treatment with DOX but without viral infection (**Fig. 6D**). Together, these data demonstrate that 3 days of FGF9 overexpression alone does not significantly alter IFN signaling in the conducting airway epithelium. Instead, high FGF9 signaling sensitizes the airway epithelial cells to induce a rapid, dramatic IFN signature, especially type I IFN, in conducting airway epithelial cells.

**Figure 6.**
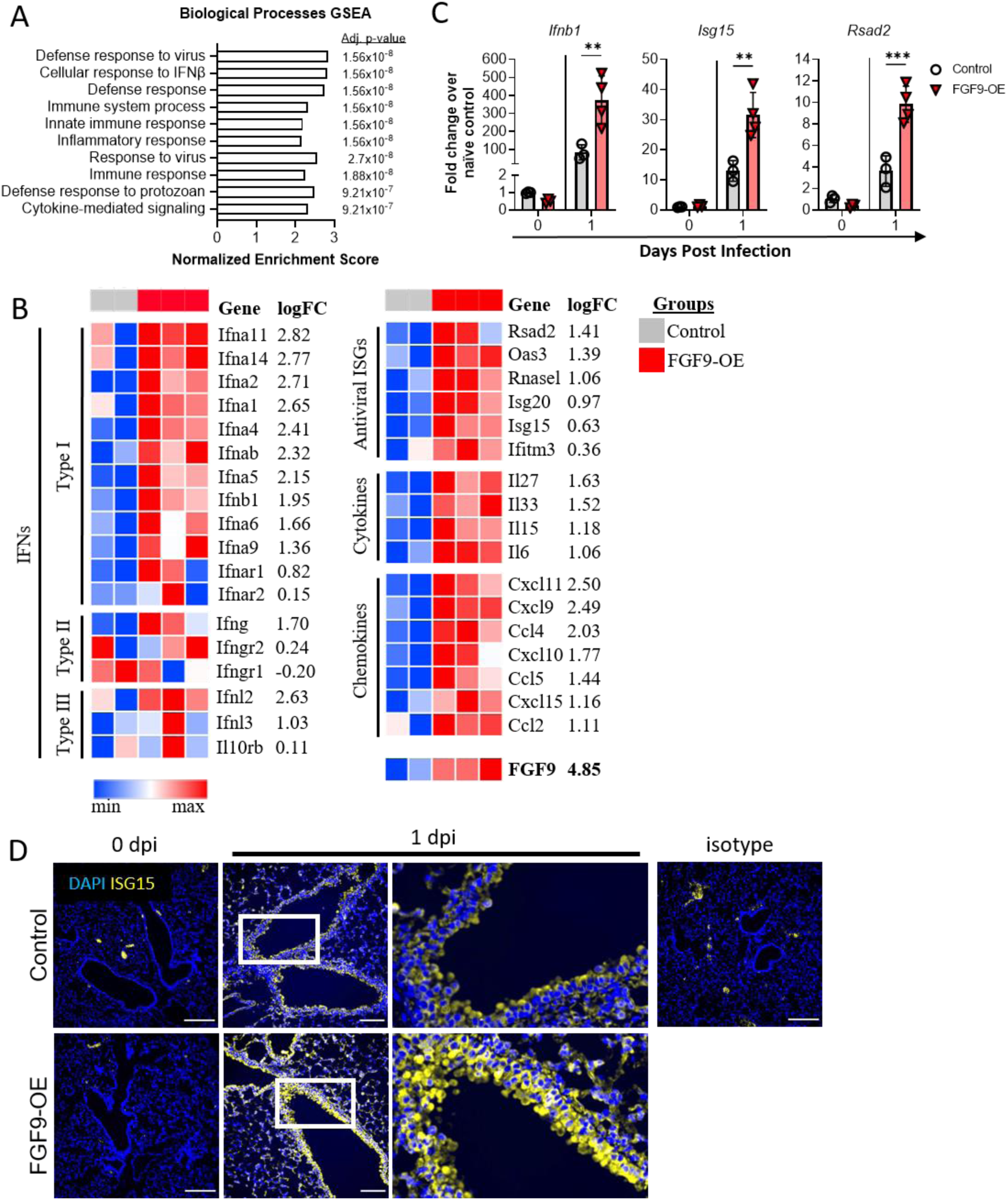
RNA sequencing revealed an elevated IFN response in FGF9-OE airway epithelial cells at 1 dpi. (A-D) FGF9-OE and control mice were administered DOX beginning on d-3 and infected on d0 with 6×10^4^ PFU WSN i.n. (A-C) RNA extracted from sorted airway epithelial cells (CD45^−^ CD326^+^ CD24^+^) at 1 dpi was analyzed by bulk RNA sequencing. (A) Top 10 most significant positively-enriched pathways, normalized enrichment scores, and adjusted p-values from GSEA analysis of Gene Ontology Biological Processes terms comparing control and FGF9-OE airway epithelial cells at 1 dpi. (B) Selected representative genes and log fold change from GSEA analysis in FGF9-OE airway epithelial cells at 1 dpi. (C) RT-qPCR validation of *Ifnb1, Isg15,* and *Rsad2* expression in sorted FGF9-OE or control airway epithelial cells at 0 and 1 dpi. Gene expression was normalized to *Gapdh* and graphed as fold change over control (0 dpi) using the 2^−ΔΔCt^ method. Data are represented as mean ± SEM and analyzed using unpaired student’s *t* test (**, p < 0.01; ***, p < 0.001). (D) Representative images of lung sections from control or FGF9-OE mice at 0 and 1 dpi stained with anti-ISG15 polyclonal sera (yellow) and analyzed by immunofluorescence microscopy (blue = DAPI, scale bars = 100 µm).

### FGF9 overexpression results in a shift in IAV tropism from the airway epithelium to the alveolar epithelium at 1 dpi

Since FGF9-OE airway epithelial cells have increased expression of ISGs, such as ISG15 and viperin, as early as 1 dpi, we wanted to investigate how this surplus of antiviral genes might affect the cellular tropism of WSN. At 1 dpi, we utilized multiplex fluorescent RNA-ISH staining of lung sections to visualize infected cells. In the control lungs, IAV^+^ cells were predominantly located in the conducting airway epithelium, with approximately 45% of airways infected and few IAV^+^ cells in the alveolar space (**Fig. 7A-C**). In stark contrast, we identified extremely low levels of infection of the conducting airway epithelial cells in the FGF9-OE mice with approximately 14% of airways infected. Instead, we observed a dramatic shift in tropism to the alveolar cells at this early time point. We propose that FGF9 overexpression in the airway epithelium protects the airway from IAV infection, likely by priming the cells to rapidly express high levels of IFNs and ISGs upon infection; however, this phenomenon precipitates an indirect consequence—rapid and robust infection of the alveolar cells, resulting in enhanced alveolar inflammation and exacerbated susceptibility to IAV disease.

**Figure 7.**
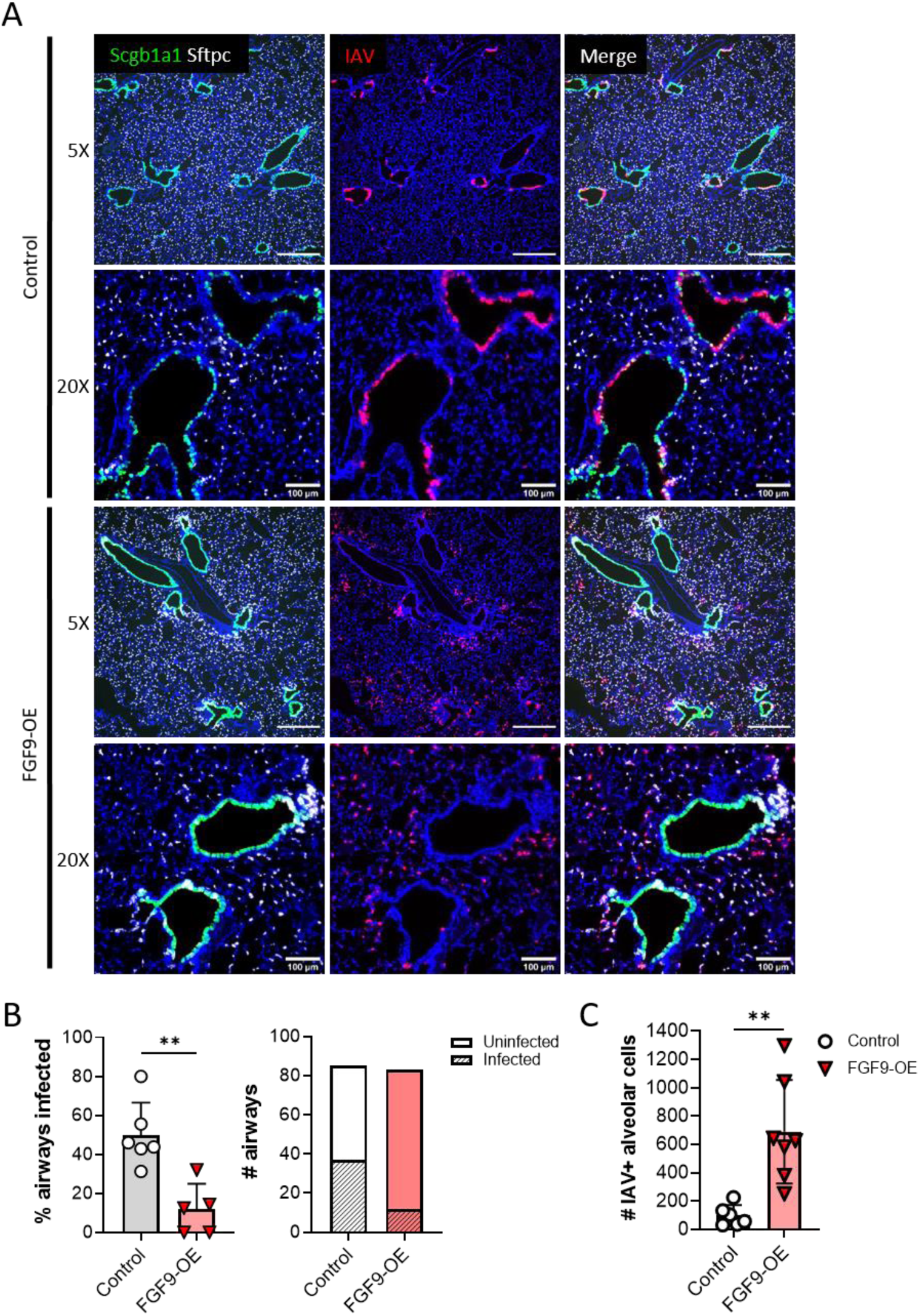
IAV tropism is shifted from the airway epithelium to the alveolar epithelium in FGF9-OE lungs at 1 dpi. FGF9-OE and control mice were administered DOX beginning on d-3 and infected on d0 with 6×10^4^ PFU WSN i.n. (A-C). (A) Representative images of multiplex fluorescent RNA-ISH depicting club cells (*Scgb1a1*, green), AT2 cells (*Sftpc*, white), and infected cells (*Np*, red) (blue = DAPI). Images taken with 5X objective (scale bars = 500 µm) and 20X objective (scale bars = 100µm). (B) The number of infected and uninfected airways was quantified from multiple FGF9-OE and control lung slides at 1 dpi, with the total percentage of infected airways per lung section (left) and total number of infected and uninfected cells (right). Infection was defined as any airway with >1 IAV+ cell. (C) The number of IAV+ cells in the alveolar space was quantified from blinded 5X images from FGF9-OE and control lung slides at 1 dpi. Data are represented as mean ± SEM and analyzed using unpaired student’s *t* test (**, p < 0.01).

## Discussion

FGFs play diverse roles in controlling IAV replication and the immune response to infection. In the current study, we found that FGF9 overexpression in club cells increased susceptibility to respiratory infection. It has been well-characterized that increased rapid infection of alveolar cells by influenza viruses worsens the severity of acute lung injury and acute respiratory distress syndrome both in humans and animal models^18, 33–37^. We found that the FGF9-OE airway epithelium had a marked reduction in IAV infection at 1 dpi, but this was accompanied by an increase of infection in the alveoli. This accelerated infection of the alveolar space could be driven by multiple mechanisms. First, the inability to infect the airway epithelium may inadvertently direct virus into the alveolar space. Alternatively, FGF9 signaling may directly disrupt the barrier function of the conducting airway epithelium, allowing virus to leak into the alveolar space during early infection. Finally, it is possible that FGF9 secreted from club cells signals directly to alveolar epithelial cells and promotes their infection. It is noteworthy that intratracheal administration of FGF7 (KGF) has been shown to increase both the proliferation of AT2 cells and their susceptibility to IAV^18^. In our FGF9-OE mouse model, FGF9 could be similarly acting on AT2 cells and promoting infection. Future work will assess whether AT2-driven FGF9 overexpression will similarly increase alveolar cell infection or instead protect the alveolar epithelium from infection in a similar manner as seen in the FGF9-OE airway epithelium.

The molecular mechanism underlying how club cell-driven FGF9 overexpression might promote a type I IFN signature upon infection remains unclear. Within the current literature, FGFs commonly inhibit or suppress the functions of IFNs in both disease and development contexts^38–40^. Recently, multiple studies have demonstrated that FGF-driven antagonism of IFN signaling can promote viral pathogenesis. For example, FGF7 treatment of keratinocytes enhanced herpes simplex virus-1, lymphochoriomeningitis virus, and Zika virus infection by inhibiting ISG expression^41^. Additionally, Zika virus infection of human fetal astrocytes elevated expression of FGF2, which suppressed IFN signaling and facilitated Zika virus infection and spread^42^. Thus, our observation of an increased IFN signature in FGF9-OE conducting airway epithelial cells unveils a potentially unique function of FGF9 signaling in the lung. FGF9 activates FGFR3 in the respiratory epithelium both during organogenesis and development of lung adenocarcinoma, whereas FGF7 and FGF10 signal through FGFR2b in the airway epithelium^12, 43, 44^. These studies also reveal that FGF signaling through FGFR3 or FGFR2b activates different signaling pathways, which could explain how FGF9 primes cells to have an increased IFN response during infection rather than inhibit IFN signaling and ISG expression. Additionally, FGF9 may instead signal on nearby fibroblasts or resident immune cells which in turn promote an elevated IFN response in the airway epithelial cells during infection. FGF9 subfamily member FGF16 was found to be antiviral against VSV in a secretome screen, and this antiviral effect was independent of IFN signaling^22^. However, the addition of IFN provided further antiviral activity against VSV in this model, demonstrating that FGF9 subfamily members may have alternative methods of providing antiviral protection in spite of or in conjunction with IFN signaling.

FGF2 can indirectly recruit neutrophils to the lung during IAV infection, thereby promoting clearance of the virus and protecting the mice from severe disease^19^. Our study shows that FGF9 overexpression similarly caused the recruitment of neutrophils, monocytes, and other innate immune cells to the lungs in conjunction with high cytokine and chemokine expression as early as 1 dpi. While it is possible that FGF9 has chemotactic properties in our IAV infection model similar to FGF2, it could be that FGF9 signaling in the lungs partially drives increased cytokine and chemokine expression resulting in the precocious recruitment of immune infiltrates, which in turn express high levels of cytokines and chemokines that we detect at 1 dpi. Indeed, our multiplex cytokine analysis revealed a modest increase in CCL2, CCL4, CXCL9, and CXCL10 in the lungs after 3 days of FGF9 overexpression.

Analysis of IAV tropism in FGF9-OE lungs at 1 dpi revealed a dramatic shift of infection into the alveolar space, which supported the severe alveolitis and alveolar edema we observed in FGF9-OE lungs at later time points. Since respiratory epithelial cells and alveolar macrophages are often the initial sources of cytokine and chemokine expression during the early stages of IAV infection, we speculate that the increased infection of alveolar cells in FGF9-OE lungs at 1 dpi provoked a surplus of cytokine and chemokine expression from both alveolar epithelial cells and alveolar macrophages.

During lung development, FGF7 and FGF10 stimulate epithelial cell proliferation and distal alveolar development, respectively, and are able to promote lung repair of the adult lung after acute lung injury^13, 14, 20, 45, 46^. Interestingly, we observe *Sftpc* expression in club cells in the FGF9-OE airway in uninfected lungs and at 1 dpi, suggesting a possible expansion of *Scgb1a1/Sftpc* (CC10/SPC) dual-positive bronchioalveolar stem cells (BASCs), which are a rare progenitor cell population normally found in healthy bronchoalveolar duct junctions in mice (**Fig. 7A, Supp. Fig. 4**). A recent study found that BASCs are able to repopulate both the airway and alveolar epithelium during recovery from IAV infection^47^, underscoring the possibility that increased highly-regenerative stem cell populations in FGF9-OE lungs may promote more rapid or largescale repair of the epithelium post infection. Future work will assess if FGF9 overexpression after IAV infection is cleared from the lungs will promote beneficial post-infection lung repair by increasing the number of BASCs in the regenerating lung.

In conclusion, we found that FGF9 overexpression in club cells in the airway epithelium locally protects the airway from early IAV infection; however, the global consequences were dire. FGF9-OE mice succumbed to severe disease, characterized by accelerated infection of alveolar cells and exacerbated inflammation. FGFs function in cellular maintenance and post-injury repair in many tissues and organs. Most treatments for viral infections focus on limiting viral replication, but understanding the contributions of FGFs and other growth factors to viral pathogenesis or protection will likely lead to better therapeutic options that optimally limit immunopathology during respiratory infection or promote tissue repair after infection.

## Materials and Methods

### Ethics Statement

Experiments were approved and performed in accordance with the recommendations in the Guide for the Care and Use of Laboratory Animals of the National Institutes of Health^48^. The protocols were approved by the Institutional Animal Care and Use Committee at the Washington University School of Medicine in St. Louis (Assurance number A-3381-01).

### Establishment of transgenic mice and doxycycline administration

The TRE-Fgf9-IRES-eGfp mice have been previously described^10^. Briefly, *Fgf9* cDNA was cloned upstream of the IRES-eGfp-SV40 polyA cassette (pIRES2-eGfp vector, Clontech) in the pTRE2 vector (Clontech)^49^. The TRE-Fgf9-IRES-eGfp transgenic mice were mated to Scgb1a1 (Ccsp)-rtTA mice^24^. All mouse strains were bred and maintained on the FVB/NJ background in our mouse colony before being genotyped and transferred to the animal biosafety level 2 laboratory for infection experiments. Mice were genotyped with PCR primers specific for Scgb1a1-rtTA (Fwd primer 5’-ACT GCC CAT TGC CCA AAC AC-3’ and Rev primer 5’-AAA ATC TTG CCA GCT TTC CCC-3’) and TRE-Fgf9 (Fwd primer 5’-GGC GTG TAC GGT GGG AGG CCT ATA TAA GC-3’ and Rev primer 5’-TAG CTC CCA ACT TCA CCT AAG GGA GCC ATC-3’). Doxycycline chow (200mg/kg) (Fisher Scientific) was used to activate the TRE transgene 3 days before viral infection, unless otherwise specified.

### Virus studies in animals

All influenza A and Sendai virus infection studies were performed in an animal biosafety level 2 laboratory and followed guidelines established by the Environmental Health and Safety Committee at Washington University School of Medicine. The principles of Good Laboratory Animal Care were adhered to in strict accordance with NIH recommendations. Every effort was made to minimize animal suffering.

Influenza A virus A/WSN/33 (WSN) was provided by Adolfo García-Sastre (Icahn School of Medicine, Mount Sinai, New York City, NY), and IAV A/PR/8/34 (PR8) was provided by Jacco Boon (Washington University School of Medicine, St. Louis, MO). Sendai virus SeV-52 was grown and characterized in the López Lab. At 8-10 weeks of age, mice were inoculated intranasally in the left nostril with 6×10^4^ PFU of WSN or 5 PFU of PR8 in a volume of 25 μl PBS or 1×10^4^ PFU SeV-52 in a volume of 40 µl PBS. Experiments included both male and female mice. For survival studies, weight loss and survival were measured daily for 14 days and mice were euthanized by carbon dioxide inhalation if they lost more than 30% of their initial body weight or were significantly moribund. At the termination of all other experiments, mice were sedated by isoflurane inhalation and euthanized via intracardiac perfusion of PBS.

### Tissue viral burden

Organs were harvested in 1 ml PBS and homogenized with 1.0 mm diameter zirconia-silica beads (BioSpec Products) with 1 pulse of 3,000 rpm for 30 sec with a MagNA Lyser (Roche) prior to plaque assay on MDCK cells. In brief, 200 µl of serial dilutions of organ homogenates in 1X DMEM (Corning) containing 0.25% BSA and 4 μg/ml N-acetylated trypsin were added to MDCK cells (5×10^5^ cells for 6 well plates) and incubated for 1 h at 37°C with rocking every 15 min. Virus was then aspirated and an MEM-agar overlay was added to the cells and incubated for approximately 72 h at 37°C. Plates were then fixed with 1% formaldehyde (for at least 30 min at room temperature) and agar plugs were removed. Plaques were visualized using a 1% crystal violet solution and counted by hand.

### Lung histology and immunofluorescence

Left lungs were inflated, fixed in 10% buffered formalin for 24-48 h, and dehydrated using 30%, 50%, and 70% ethanol. Lungs were subsequently paraffin-embedded and sectioned at 5 μm. For histological analysis, sections were deparaffinized with xylene, stained with hematoxylin and eosin (H&E), and evaluated by light microscopy using a Zeiss Axio Scan.Z1 slide scanner. H&E-stained lung sections were blinded and scored on a scale of 0 to 3 for peribronchiolitis, squamous epithelial cell metaplasia, alveolitis, and airway epithelial denudation. Percentage of airways and the area of the lung affected for each type of pathologic finding was determined. The raw percentage of affected area was multiplied by the intensity score to acquire the graphed weighted score^50^.

For immunofluorescence, 5 μm sections were deparaffinized with xylene, boiled between 98°-102°C for 15 min with Antigen Unmasking Solution (Vector Laboratories), and permeabilized for 10 min at room temperature with PBS/2% Triton TX-100/1% sodium citrate. Sections were blocked using PNB buffer (0.5% PNB powder in PBS), antibody and DAPI (4′,6-diamidino-2-phenylindole) stained, and visualized by fluorescence microscopy. The following primary antibodies were used: anti-ISG15 polyclonal rabbit sera (1:250)^51^, anti-SCGB1A1 goat polyclonal antibody Sc9772 (1:200) (Santa Cruz). Embedding and sectioning were performed by the Musculoskeletal Histology and Morphometry Core at Washington University School of Medicine. All samples were visualized using a Zeiss Axio Imager Z2 fluorescence microscope with ApoTome2 and processed with Zen Pro software.

### Multiplex fluorescence RNA *in situ* hybridization (RNA-ISH)

Multiplex fluorescence RNA-ISH was performed using RNAscope Multiplex Fluorescent Reagent Kit v2 (Advanced Cell Diagnostics, Inc., Newark, NJ, USA) according to the manufacturer’s instructions. Briefly, left lungs were inflated, fixed in 10% buffered formalin, and paraffin embedded. 5 μm sections were deparaffinized, digested with protease, and hybridized for 2 h at 40°C with the following RNAscope target RNA-specific probes: V-Influenza-H1N1-H5N1-NP (Cat. No. 436221), Mm-Sftpc-C2 (Cat. No. 314101-C2), and Mm-Scgb1a1-C3 (Cat. No. 420351-C3). Preamplifier, amplifier, HRP-labeled oligos, and TSA plus Cyanine3, Cyanine5, and fluorescein dyes were then hybridized sequentially at 40°C. The nuclei were stained with DAPI, and images were acquired as described above. Infection of airways was quantified by counting the number of infected (>1 IAV+ cell in the airway epithelium) or uninfected airways in multiple lung sections from 4 FGF9-OE and 4 control mice. Infection of alveoli was quantified by counting the number of IAV+ cells not associated with the airway epithelium or endothelium in 3-4 5X images spanning entire lung lobe sections from 4 FGF9-OE and 3 control mice.

### Cytokine and chemokine multiplex analysis

Whole lungs were collected in 1 ml PBS with 0.1% BSA at 0, 1, 3, and 5 dpi and homogenized with 1.0 mm diameter zirconia-silica beads (BioSpec Products) using a MagNA Lyser (Roche). Cytokines and chemokines in the homogenates were quantified using Luminex technology with a MILLIPLEX Map Mouse Cytokine/Chemokine Magnetic Bead Panel 25-plex Immunology Multiplex Assay (Cat. No. MCYTOMAG-70K-PMX, Millipore) according to the manufacturer’s instructions, with modifications recommended for “sticky” samples in the Millipore “tips and tricks” brochure (version 1.0 2017-01180). In brief, these modifications included pelleting tissue debris with a 5 min centrifugation step (600 × g) before adding samples to the 96-well assay plate and running the plate in 1X wash buffer instead of sheath fluid.

### Real-time qPCR analysis of host gene expression

Lungs were collected for RNA processing by snap freezing in liquid nitrogen in tubes with 1.0 mm diameter zirconia-silica beads (BioSpec Products) and transfered to -80°C for storage. Whole lung RNA was extracted from the tissue using TRIzol Reagent and an RNeasy Mini Kit (Qiagen). The TRIzol Reagent manufacturer’s protocol was followed up until removal of the RNA-containing aqueous phase, to which an equal volume of 100% RNase-free ethanol was added. This solution was added to the RNeasy column, after which the RNeasy Mini Kit instructions were followed.

For host gene expression, 2 µl of the extracted RNA from WSN-infected or uninfected mouse lungs were analyzed by quantitative real time PCR (RT-qPCR) using the TaqMan RNA-to-Ct 1-Step Kit (Applied Biosystems). Ct values for the gene of interest and the housekeeping gene *Gapdh* (glyceraldehyde-3-phosphate dehydrogenase) were used to calculate ΔΔCt, and normalized. Fold change was calculated by 2^-ΔΔCt^ method^52^. Custom Taqman primers and probes were acquired from Integrated DNA Technologies to detect *Ifnb1*: *Ifnb1* FW (5′ GTT GAT GGA GAG GGC TGT G 3′), *Ifnb1* probe (5′ /56-FAM/CTG CGT TCC/ ZEN/ TGC TGT GCT TCT C /3IABkFQ/ 3′), *Ifnb1* RV (5′ GGC TTC CAT CAT GAA CAA CAG 3′)^53^. Standard assays from Integrated DNA Technologies were utilized to detect *Gapdh* (Mm.PT.39a.1), *Isg15* (Mm.PT.58.41476392.g), and *Rsad2* (Mm.PT.58.11280480).

### Flow cytometry and analysis

Mice were sacrificed and perfused with PBS as described above. Lung single cell suspensions were generated by incubation for 1 h at 37°C in 5 ml digestion buffer with manual shaking every 10-15 min. Digestion buffer for epithelial cell analysis consisted of RPMI (Sigma), DNase I (10 mg/ml, Sigma), 15 mM HEPES buffer (Corning), and 10% fetal bovine serum (FBS) (BioWest). Digestion buffer for immune cell analysis additionally included type IV collagenase (2.5 mg/ml, Sigma). Digested tissues were passed through a 70-µm cell strainer and washed once with 40 ml PBS containing 5% FBS and treated with red blood cell lysing buffer (Sigma). The number of viable cells was quantified by trypan blue staining.

The single cell suspension was transferred to a 96-well v-bottom plate, and Fcγ receptors were blocked with anti-mouse CD16/CD32 (Clone 93; BioLegend) for 15 min at 4°C, followed by surface staining in PBS containing 5% FBS for 1 h at 4°C. All antibodies were diluted 1:200 and are from BioLegend unless otherwise specified: Fixable Viability Dye eFluor 506 (1:500, eBioscience); anti-CD45.1 Pacific Blue (A20); anti-I-A/I-E PerCP-Cy5.5 (peridinin chlorophyll protein-Cyanine5.5) (M5/114.15.2); anti-CD11b Brilliant Violet 605 (M1/70); anti-Siglec F APC-Cy7 (allophycocyanin-Cyanine7) (E50-2440); anti-Ly6C Alexa Fluor 700 (HK1.4); anti-CD11c APC (allophycocyanin) (N418); anti-Ly6G PE-Cy7 (phycoerythrin-Cyanine7) (1A8); anti-CD103 PE (phycoerythrin) (2E7); anti-CD45 Alex Fluor 700 (30-F11); anti-CD326 PE (phycoerythrin) (G8.8); anti-CD24 APC (allophycocyanin) (M1/69).

After staining, cells were washed and fixed at 4°C for 10 min in 4% paraformaldehyde (Electron Microscopy Sciences). The fixed cells were washed and resuspended in PBS containing 5% FBS. Cells were processed on a LSR Fortessa (Becton Dickinson) flow cytometer managed by the Flow Cytometry & Fluorescence Activated Cell Sorting Core at Washington University and analyzed using BD FACSDiva and FlowJo V10 software (Tree Star Inc.).

### Bulk RNA sequencing from sorted cells

Whole lung lysates were processed for flow cytometry as described above using epithelial cell digestion buffer. Whole lungs were combined from 2 mice for each single sample. Single cell suspensions were incubated with anti-mouse CD16/CD32 (Clone 93; BioLegend) for 15 min at 4°C and then surface stained in PBS containing 5% FBS for 1 h at 4°C. All antibodies were diluted 1:200 and are from BioLegend: anti-CD45 Alex Fluor 700 (30-F11), anti-CD326 (EpCAM) PE (phycoerythrin) (G8.8), anti-CD24 APC (allophycocyanin) (M1/69).

After staining, cells were washed in PBS containing 5% FBS and CD45^−^ EpCAM^+^ CD24^+^ cells were sorted using a Sony Cell Sorter SH800S and Cell Sorter Software V2.1.5 (Sony Biotechnology, Sony Biotechnology). RNA was extracted as described above for RT-qPCR analysis. 3 FGF9-OE and 2 control RNA samples were submitted to the Genome Technology Access Center at the McDonnell Genome Institute at Washington University School of Medicine for the following analyses. Total RNA integrity was determined using Agilent Bioanalyzer. Library preparation was performed with 10 ng of total RNA with a Bioanalyzer RIN score greater than 8.0. ds-cDNA and was prepared using the SMARTer Ultra Low RNA kit for Illumina Sequencing (Takara-Clontech) per manufacturer’s protocol. cDNA was fragmented, blunt ended, had an A base added to the 3’ ends, and then had Illumina sequencing adapters ligated to the ends. Ligated fragments were then amplified for 12-15 cycles using primers incorporating unique dual index tags. Fragments were sequenced on an Illumina NovaSeq-6000 using paired end reads extending 150 bases. High quality gene reads were mapped to the *Mus musculus* genome and differential gene expression of normalized counts per million was analyzed using the Phantasus platform (https://artyomovlab.wustl.edu/phantasus/) developed by the Sergushichev team at ITMO university. Briefly, gene expression values were *Log 2* and *quantile* normalized and the top 12,000 most expressed genes were subjected to differential gene expression analysis using the *limma* R package. The Gene Ontology Biological Processes database was used to for pathway enrichment analysis.

### Statistical Analysis

All data were analyzed using the Prism software, version 9 (GraphPad), as detailed in the figure legends using Mantel-Cox test, unpaired, two-tailed student’s *t* test, or two-way ANOVA with Tukey’s post-test for multiple comparisons statistical tests. Statistical outliers (p<0.05) were removed by Grubb’s test for multiplex cytokine/chemokine analysis. All error bars indicate SEM; if error bars are not visible, then they are shorter than the height of the symbol. Asterisks indicate statistical significance, with only relevant comparisons shown (*, *p* < 0.05; **, *p* < 0.01; ***, *p* < 0.001; ****, *p* < 0.0001).

## Acknowledgements

This study was supported by grants from the NIH (R01 AI080172 to D.J.L., 5T32 CA009547 to Y.-C.P. and L.E.F., 5T32 AI007163 and F31 AI 14999 to M.C.L., R01 AI134862 and R01 AI137062 to I.A.C. and C.B.L., and R01 HL111190 and R01 HL154747 to D.M.O.) We also acknowledge support from the Washington University Rheumatic Diseases Research Resource-based Center (P30 AR073752) for genome engineering and sequencing.

We acknowledge that microscopy imaging was performed in part through the use of the Washington University Center for Cellular Imaging (WUCCI), which was supported by the Washington University School of Medicine, The Children’s Discovery Institute of Washington University, St. Louis Children’s Hospital (CDI-CORE-2015-505), Washington University Rheumatic Diseases Research Resource-based Center (P30 AR073752), and by the Foundation for Barnes-Jewish Hospital (3770). We acknowledge that all processing of tissues for histology and H&E staining was performed by the Washington University Musculoskeletal Research Center, which received funding from the NIH (P30 AR057235). We thank the Genome Technology Access Center at the McDonnell Genome Institute at Washington University School of Medicine for help with RNA-seq analysis. The Center is partially supported by NCI Cancer Center Support Grant #P30 CA91842 awarded to the Siteman Cancer Center and by ICTS/CTSA Grant# UL1TR002345 from the National Center for Research Resources (NCRR), a component of the National Institutes of Health (NIH), and by NIH Roadmap for Medical Research. This publication is solely the responsibility of the authors and does not necessarily represent the official view of NCRR or NIH.

The funders had no role in study design, data collection and analysis, decision to publish, or preparation of the manuscript.

---

**Supplemental Figure 1.**
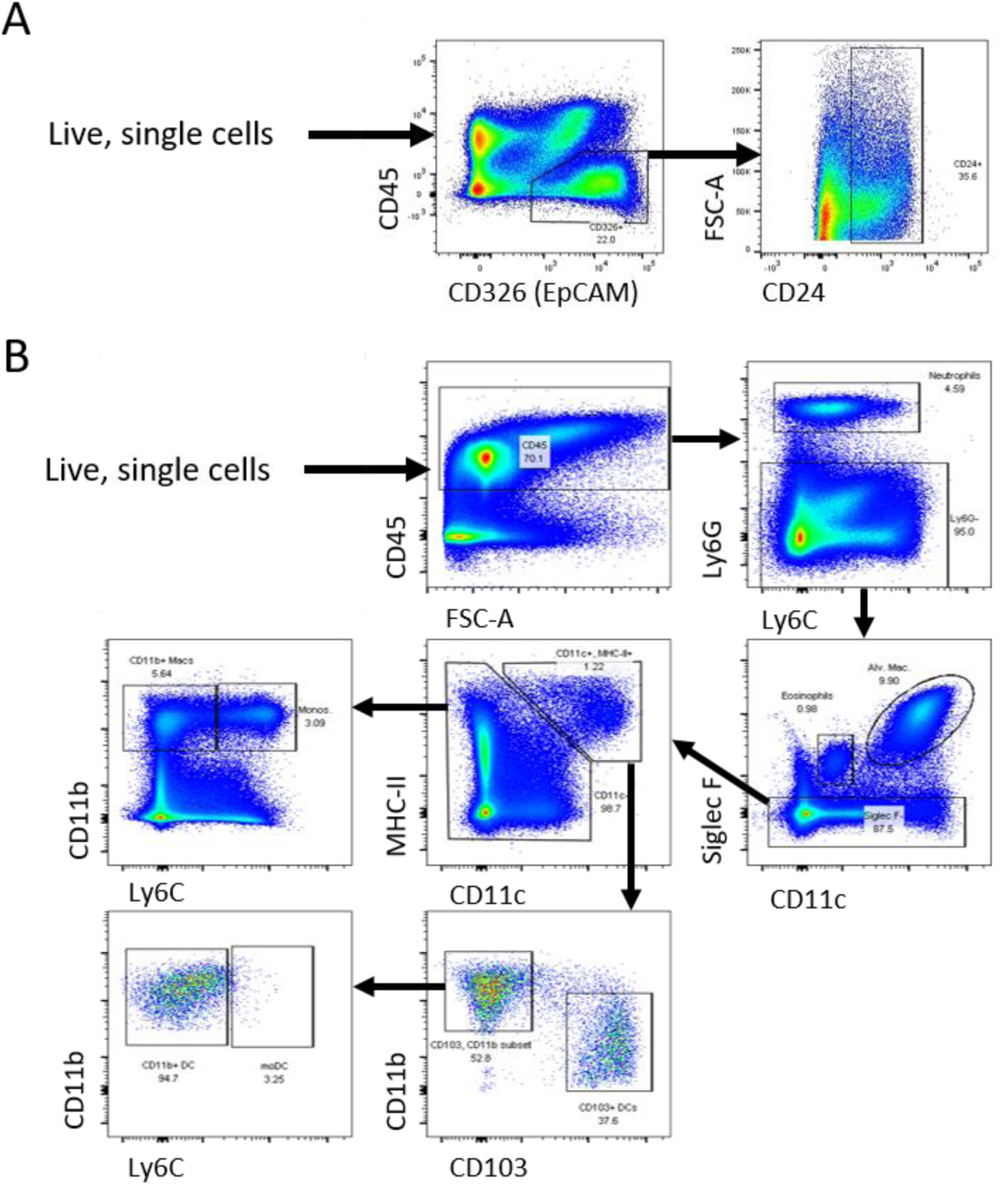
Gating strategies for isolating respiratory epithelial cells and innate immune cells by flow cytometry. (A-B) Representative flow cytometry plots depicting the gating strategy for epithelial cell subpopulations (A) and innate immune cell (B) subpopulations isolated from control mice lungs.

**Supplemental Figure 2.**
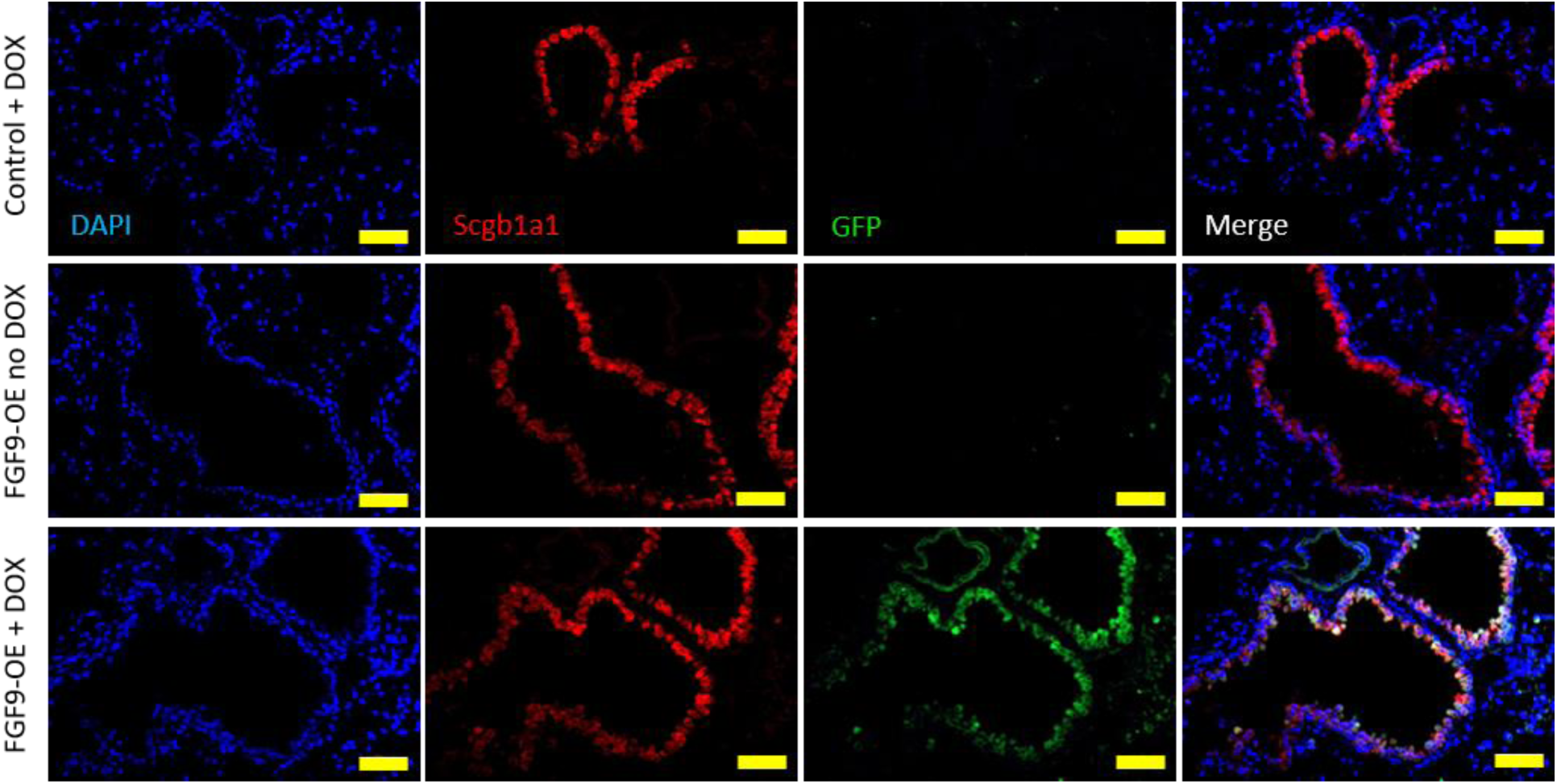
FGF9-OE club cells required doxycycline to express eGFP from the TRE-Fgf9-IRES-eGfp cassette. Representative images of lung sections from control mice or FGF9-OE mice administered DOX for 3 days and FGF9-OE mice without DOX staining for DAPI (blue) and Scgb1a1 (red) and endogenous eGFP expression. Scale bars = 50 μm.

**Supplemental Figure 3.**
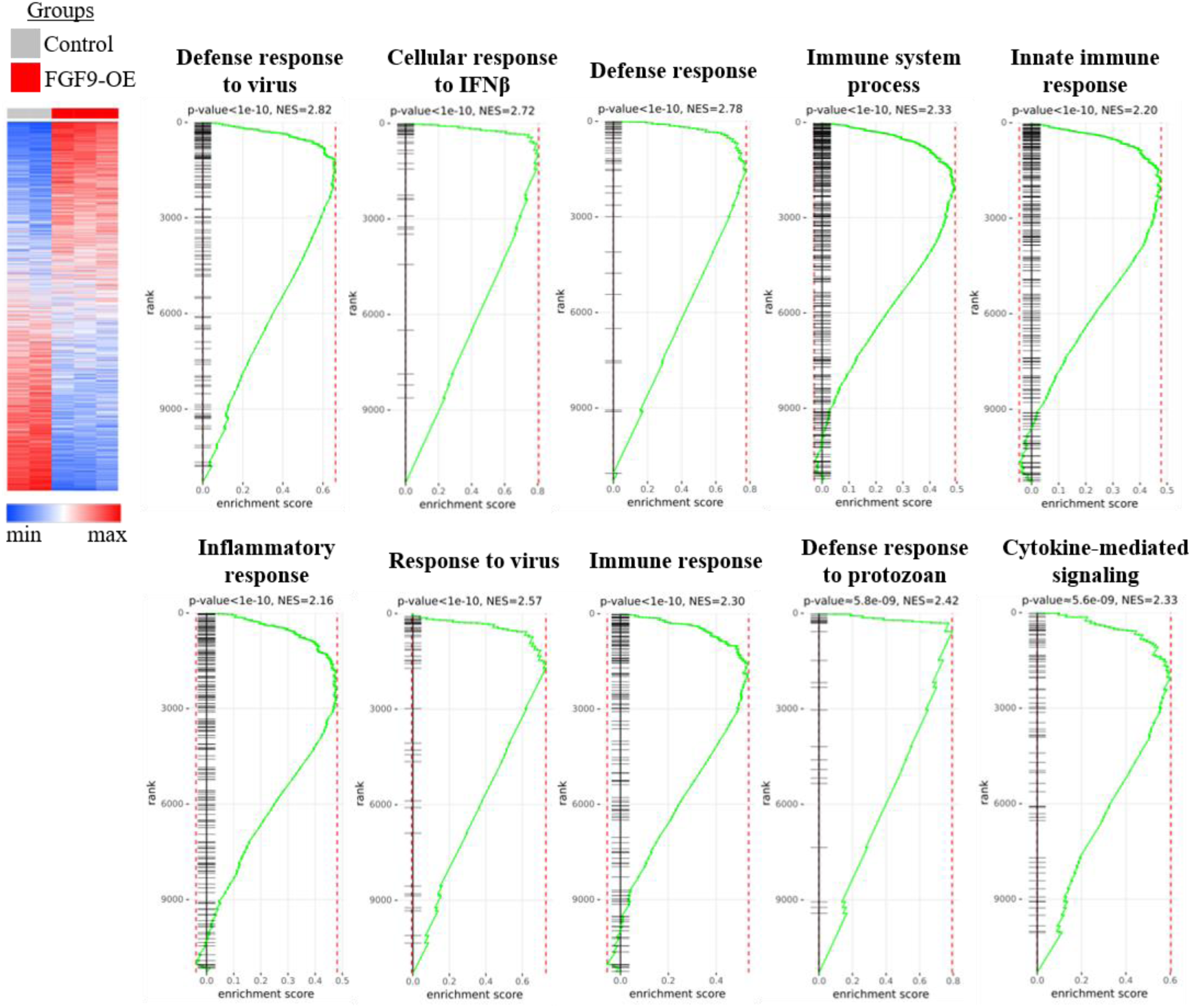
GSEA plots of the most positively-enriched pathways in FGF9-OE airway epithelial cells at 1 dpi. GSEA plots of the top ten most significant (by adjusted p-value) positively-enriched pathways found upregulated in the FGF9-OE airway epithelial cells as compared to control airway epithelial cells at 1 dpi.

**Supplemental Table 1.**
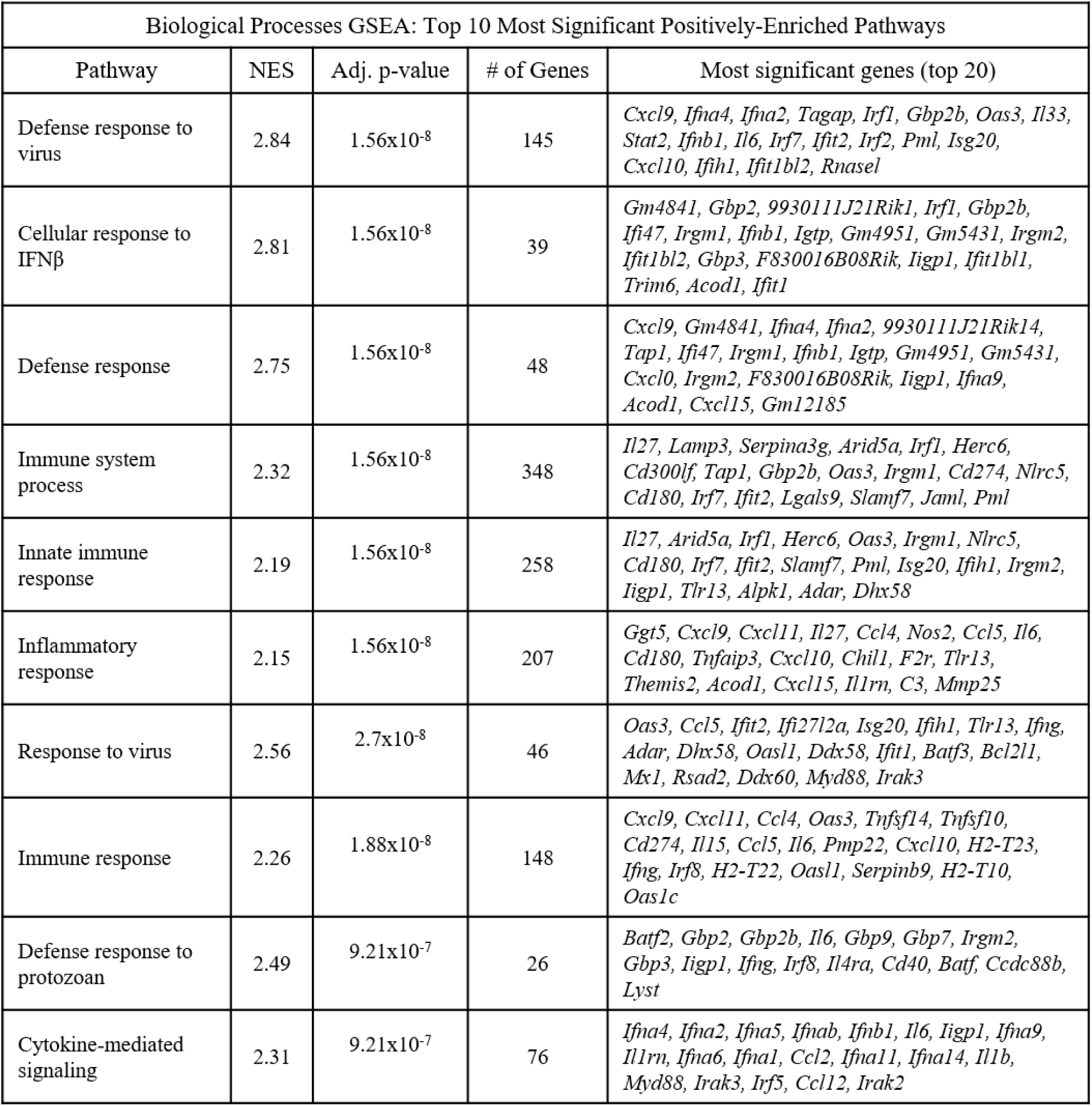
List of genes in the most significant positively-enriched pathways in FGF9-OE airway epithelial cells at 1 dpi. Normalized enrichment scores (NES), number of genes, and the top 20 most significant genes in each pathway of the top ten most significant (by adjusted p-value) positively-enriched pathways upregulated in the FGF9-OE airway epithelial cells compared to control airway epithelial cells at 1 dpi.

**Supplemental Figure 4.**
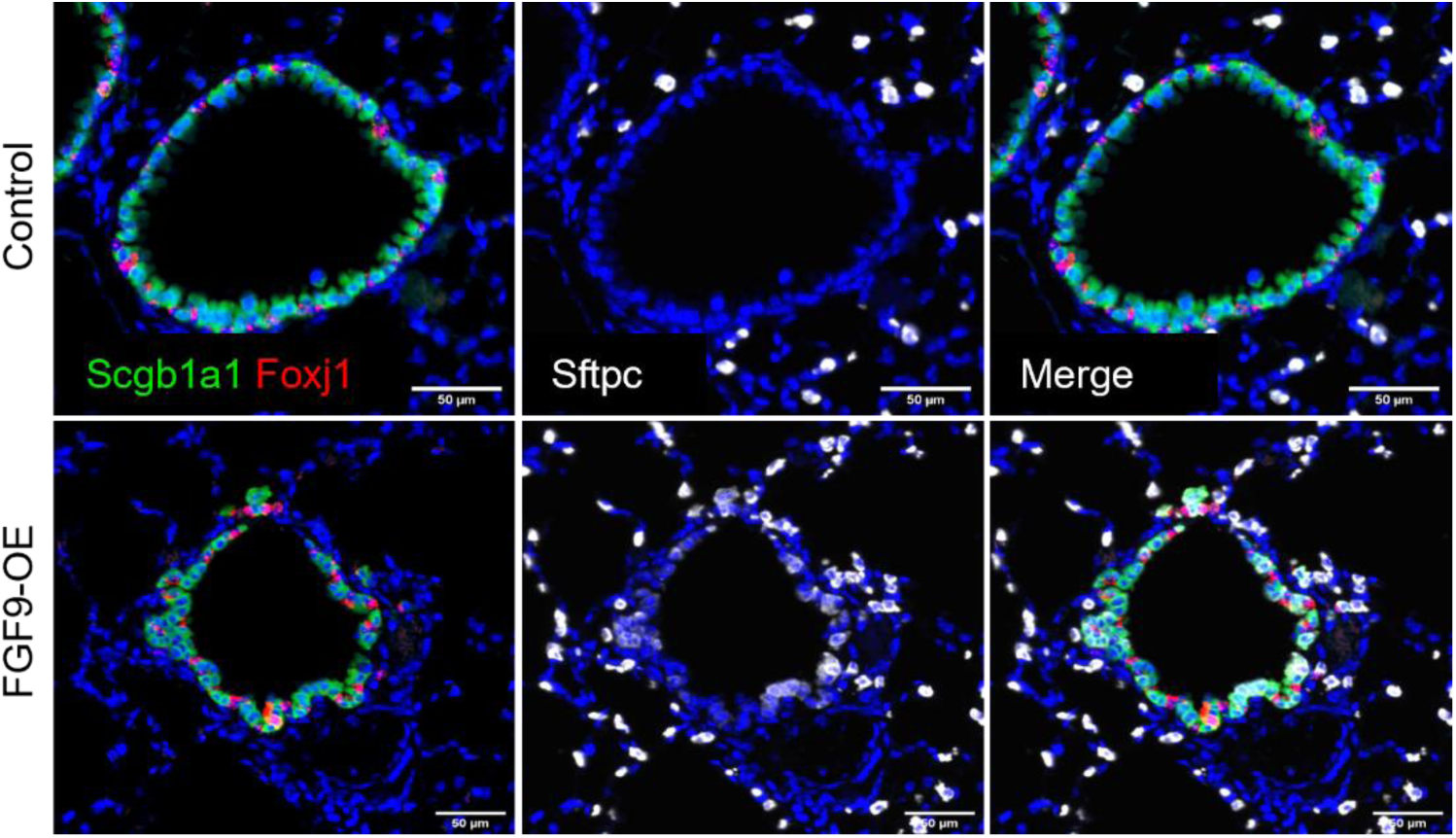
S*f*tpc expression in FGF9-OE club cells. FGF9-OE and control mice were administered DOX beginning on d-3 and lungs were harvested on d0. Shown are representative images of multiplex fluorescent RNA-ISH showing club cells (*Scgb1a1*, green), AT2 cells (*Sftpc*, white), and ciliated cells (*Foxj1*, red) (blue = DAPI). Images taken with 20X objective (scale bars = 50µm).

